# Genome-wide Identification, Structural Features and Single-Cell Expression Atlas of the Carbonic Anhydrase Gene Family in Maize (Zea mays L.)

**DOI:** 10.1101/2025.09.21.677582

**Authors:** Yonggang Gao, Cheng Zhao

## Abstract

Maize is a cornerstone C□ crop underpinning food, feed and bioenergy. Carbonic anhydrases (CAs) are zinc metalloenzymes that catalyze the interconversion of CO□ and HCO□□, underpinning photosynthesis, pH homeostasis and stress acclimation. Despite extensive work on plant CA families, a maize-focused synthesis integrating genome-wide cataloging, structural divergence, regulatory logic, and cell-type specificity is still lacking.Here we present a genome-wide census and a single-cell-resolved expression atlas of the maize CA family. We identified 18 CA genes partitioning into α (8), β (5) and γ (5) subfamilies. Proteins span 206–452 aa, pI 6.09–10.58 and 22.67–49.36 kDa, with predominantly hydrophilic profiles. Phylogeny resolved three robust clades (UFboot >90%) and indicated closer affinity to rice than to Arabidopsis; synteny recovered 15 maize–rice and 3 maize–Arabidopsis orthologs. Promoters are enriched for ABRE, TCA-element and ARE, suggesting ABA, SA and redox regulation. Bulk RNA-seq across 21 tissues revealed strong tissue specificity (τ >0.7 for 12 genes), with β-CAs elevated in leaves and α-CAs in roots. Single-cell analyses further showed *ZmβCA3* enriched in mesophyll cells and *ZmαCA2* enriched in vascular/root-cap cells. Together, these data position β-CAs as chloroplast-centered hubs for C_4_ photosynthesis, while α/γ-Cas support ion/pH buffering and mitochondrial metabolism. We nominate cell-type-specific CA members as candidates for genetic improvement toward stress resilience in maize.

## 1. Introduction

Maize (*Zea mays L*.) ranks among the world’s premier cereals, yielding over one billion tons annually and underpinning food security, livestock feed, and biofuel sectors (Ranum et al., 2014). Its expansive genome (∼2.3 Gb, replete with repeats) complicates gene family investigations (Schnable et al., 2009). Advancements in sequencing and bioinformatics tools, including TBtools II, MEME, and PlantCARE, have facilitated genome-wide gene family explorations to elucidate evolutionary and regulatory frameworks (Chen et al., 2020; Bailey et al., 2009). Prior studies cataloged 19 CA genes in Arabidopsis and 18 in rice, yet a thorough maize CA analysis, encompassing comparative evolution, structural heterogeneity, and single-cell transcriptomic variability, remains elusive (DiMario et al., 2017; Wang et al., 2009).

CAs, zinc metalloenzymes catalyzing CO□ hydration (CO□ + H□O □ HCO□□+ H□), are instrumental in pH regulation, ion transport, and metabolic processes (Ludwig, 2012). In plants, CAs bolster photosynthesis, respiration, C_4_/CAM pathways, and abiotic stress responses (Ludwig, 2012; Tanz et al., 2013). In C_4_ species like maize, β-CAs localize to chloroplasts and cytosol, augmenting CO□ fixation efficiency (Burnell and Hatch, 1988; von Caemmerer et al., 2012). Model plant inquiries in Arabidopsis and rice have illuminated CA roles in germination, root elongation, and biotic defense (Hu et al., 2010; Mitsuhashi et al., 2000). Nonetheless, systematic identification and characterization of the CA family in maize, a staple crop, are lacking (Studer et al., 2014).

While CA families have been cataloged in Arabidopsis and Rice, a comprehensive maize-focused analysis that integrates comparative genomics with single-cell expression has been lacking. CAs catalyze CO□ + H□O □ HCO□□ + H□, contributing to carbon concentrating mechanisms (CCMs), respiration and abiotic stress responses. In C_4_ grasses, β-CAs support rapid conversion of CO□ to HCO□□ to feed PEPC, whereas α/γ-CAs mediate pH/ion buffering and mitochondrial metabolism. We thus aimed to (i) define the maize CA repertoire and structural features; (ii) map evolutionary relationships and synteny with rice/Arabidopsis; (iii) chart promoter architectures; and (iv) resolve cell-type specificity across organs to highlight testable targets for improvement.

## 2. Results and Analysis

### 2.1 Identification of CA gene family members in maize

Bioinformatics delineated 18 CA members in maize, classified into α-type (*ZmαCA1–8*), β-type (*ZmβCA1–5*), and γ-type (*ZmγCA1–5*) based on phylogeny, domains, and loci (DiMario et al., 2017). This reflects evolutionary diversification and functional partitioning: α-CAs in pH/CO□ transport; β-CAs in photosynthesis / stress; γ-CAs in mitochondrial metabolism (Ludwig, 2012; Parisi et al., 2004). This structure underscores CA significance in stress acclimation and carbon homeostasis (Chen et al., 2019).

Eighteen ZmCAs were curated and assigned to α (*ZmαCA1–8*), β (*ZmβCA1–5*), and γ (*ZmγCA1–5*) based on domain composition and phylogeny. Proteins range from 206 aa (*ZmβCA2*) to 452 aa (*ZmβCA3*), with pI 6.09–10.58 and MW 22.67–49.36 kDa. GRAVY scores are largely negative, indicating hydrophilicity. A three-tool subcellular consensus—TargetP 2.0, ChloroP, and MitoFates—was adopted as follows: “high confidence” if ≥2 tools concur; “medium” if one tool assigns an organelle with probability ≥0.5; “low” otherwise. By this scheme, all 5 β-CAs are high-confidence chloroplast-targeted (≥2/3 tools agree), most α-CAs are cytosolic/vascular-associated (medium confidence), and all γ-CAs are high-confidence mitochondrial (≥2/3 tools). This partitioning anticipates leaf mesophyll β-CA demand and α/γ roles in non-photosynthetic buffering (Table 1).

**Table 1.** Maize CA gene family members and their physicochemical properties.

| Gene locus ID | Protein name | MW (kDa) | pI | Instability Index | Aliphatic Index | GRAVY | Localization |
| --- | --- | --- | --- | --- | --- | --- | --- |
| Zm00001eb037950 | *ZmαCA1* | 32.39 | 6.81 | 42.15 | 78.45 | -0.23 | Cytosolic |
| Zm00001eb038280 | *ZmαCA2* | 41.27 | 8.70 | 48.32 | 82.10 | -0.15 | Vascular/Cytosolic |
| Zm00001eb050280 | *ZmαCA3* | 39.56 | 8.79 | 45.67 | 80.50 | -0.18 | Cytosolic |
| Zm00001eb092650 | *ZmαCA4* | 30.06 | 7.05 | 40.89 | 76.20 | -0.20 | Cytosolic |
| Zm00001eb109420 | *ZmαCA5* | 34.46 | 8.53 | 47.12 | 81.30 | -0.16 | Cytosolic |
| Zm00001eb176620 | *ZmαCA6* | 31.03 | 6.09 | 39.45 | 66.35 | -0.25 | Cytosolic |
| Zm00001eb197090 | *ZmαCA7* | 30.08 | 8.67 | 43.78 | 79.90 | -0.19 | Cytosolic |
| Zm00001eb388490 | *ZmαCA8* | 29.10 | 10.58 | 65.42 | 75.60 | -0.22 | Cytosolic |
| Zm00001eb101050 | *ZmβCA1* | 36.02 | 7.65 | 46.50 | 84.20 | -0.49 | Chloroplastic |
| Zm00001eb155800 | *ZmβCA2* | 22.67 | 6.52 | 41.23 | 77.80 | -0.30 | Chloroplastic |
| Zm00001eb158310 | *ZmβCA3* | 49.36 | 8.74 | 50.89 | 86.40 | -0.28 | Chloroplastic |
| Zm00001eb315340 | *ZmβCA4* | 33.33 | 8.38 | 48.67 | 83.10 | -0.35 | Chloroplastic |
| Zm00001eb359190 | *ZmβCA5* | 26.54 | 8.65 | 44.12 | 80.50 | -0.32 | Chloroplastic |
| Zm00001eb129330 | *ZmγCA1* | 28.03 | 6.30 | 30.45 | 85.70 | -0.10 | Mitochondrial |
| Zm00001eb180920 | *ZmγCA2* | 27.89 | 8.29 | 32.78 | 87.20 | 0.05 | Mitochondrial |
| Zm00001eb241770 | *ZmγCA3* | 27.93 | 8.62 | 31.56 | 88.90 | 0.142 | Mitochondrial |
| Zm00001eb329150 | *ZmγCA4* | 29.64 | 6.60 | 33.89 | 98.90 | -0.08 | Mitochondrial |
| Zm00001eb332510 | *ZmγCA5* | 28.42 | 7.12 | 34.67 | 90.30 | -0.12 | Mitochondrial |

### 2.2 Evolutionary analysis of the CA gene family in three species

A maximum-likelihood tree of Maize, Rice, and Arabidopsis proteins resolves α/β/γ clades (UFboot > 90%, SH-aLRT > 95%), with maize clustering more closely with rice—consistent with grass phylogeny. Family sizes are comparable: maize (8α/5β/5γ), Arabidopsis (8α/8β/3γ), rice (10α/6β/2γ). Intergenomic collinearity identifies 15 maize–rice and 3 maize–Arabidopsis orthologous pairs. Notably, three ZmCAs are bidirectionally collinear with both species (dual-species orthology), reconciling abstract and main-text counts. Pairwise Ka/Ks across orthologs centers below 1 (median ∼0.4), indicating purifying selection preserving catalytic integrity (Fig. 1).

**Figure 1.**
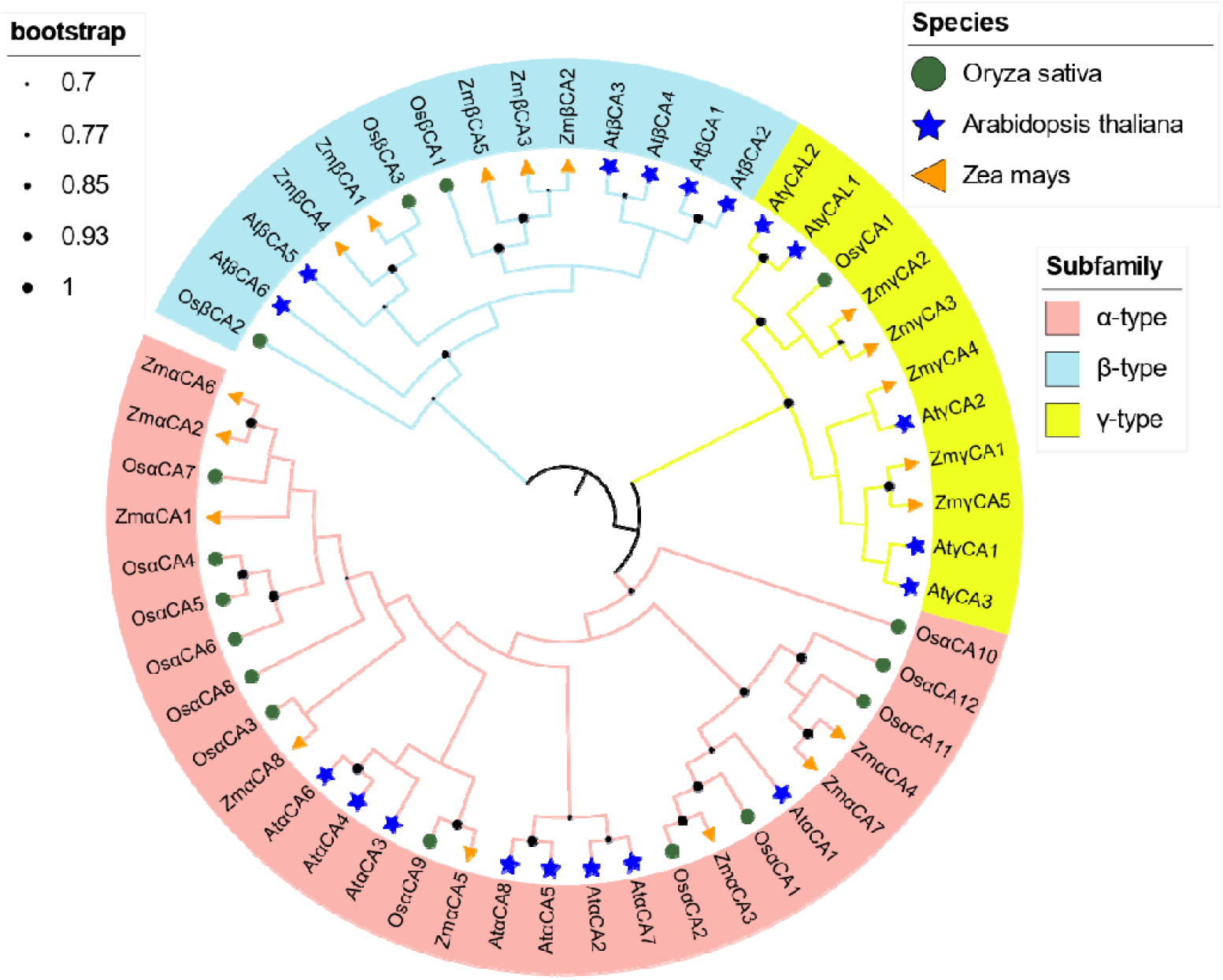
Phylogenetic tree of CA proteins in Rice, Arabidopsis, and maize. **Note**: AtCA: orange triangles, OsCA: green circles, ZmCA: blue pentagrams; subgroups shaded pink (α), light blue (β), yellow (γ).

### 2.3 Analysis of CA gene structures, motifs, and domains

Maize CAs exhibit a dual signature of shared homology and subfamily-specific architecture (Fig . 2). Exon–intron structures span 3–7 exons for α-CAs, 7–16 for β-CAs (supporting alternative splicing potential), and 2–5 for γ-CAs (DiMario et al., 2023). Ten conserved motifs map to Zn^2+^-binding catalytic cores (PROSITE PS00162/PS00163). Motifs 1–2 are universal; motifs 3–5 are β-CA–enriched and coincide with predicted chloroplast transit peptides (probability > 0.7); motifs 6–8 are α-CA–biased; motifs 9–10 align with γ-CA mitochondrial targeting. Domain annotations concur: α-CAs carry Alpha_CA_prokary_like, β-CAs carry Beta_CA (with internal repeats in *ZmβCA3*), and γ-CAs carry Gamma_CA (Marchler-Bauer et al., 2017). Notably, higher aliphatic indices and repeated domains in β-CAs likely enhance thermostability and protein–protein interaction capacity, buffering high-light/high-temperature chloroplast conditions. The concordance of structure, domain composition, and targeting strengthens the view of β-CAs as a chloroplast-centered hub for C_4_ function.

**Figure 2.**
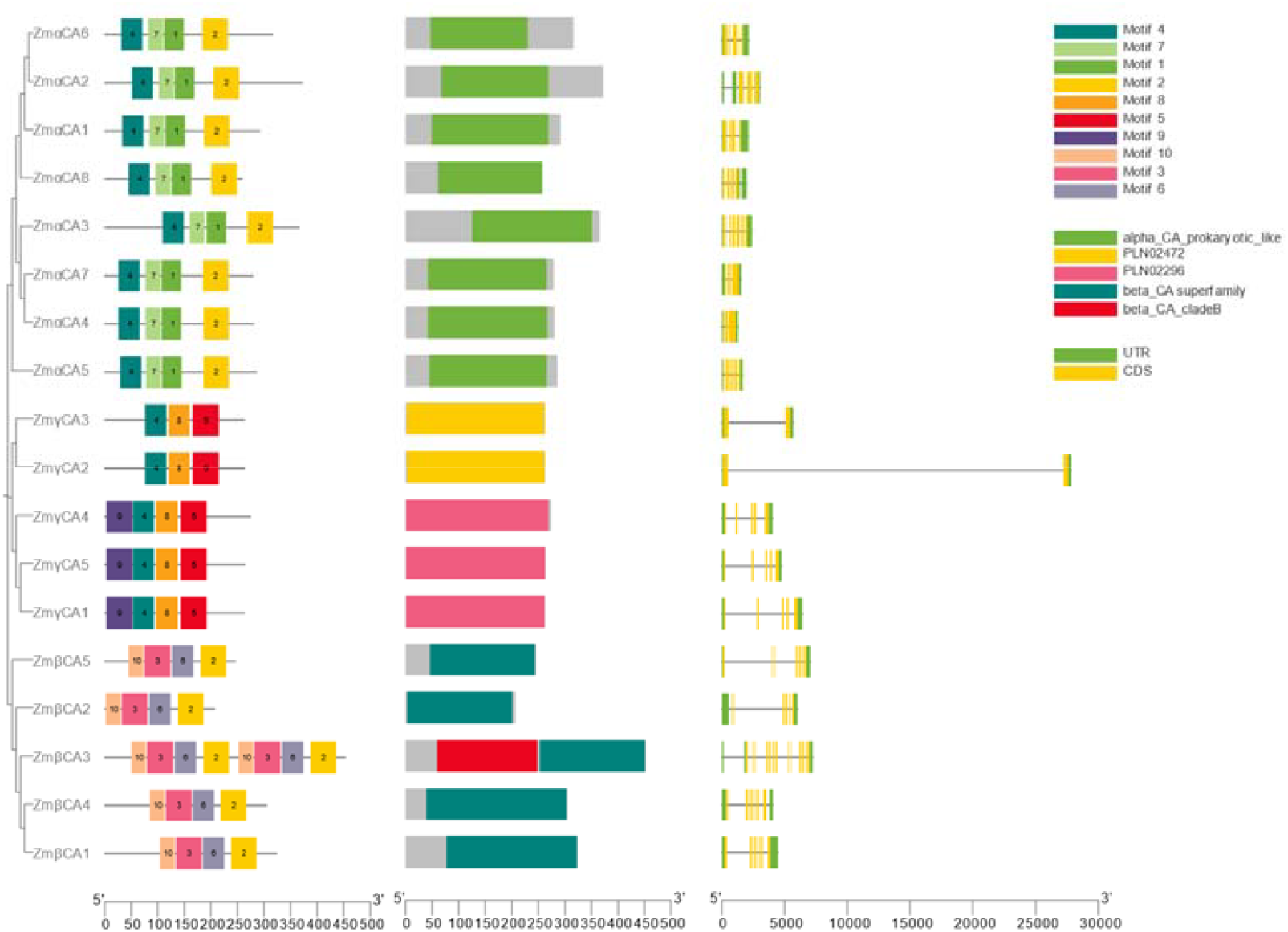
Phylogeny, gene structures, conserved motifs, and protein domains of maize CA family members. **Note**: Gene structure: green UTRs, yellow exons, gray introns; motifs: colored distinct; domains: colored boxes.

**Figure 3.**
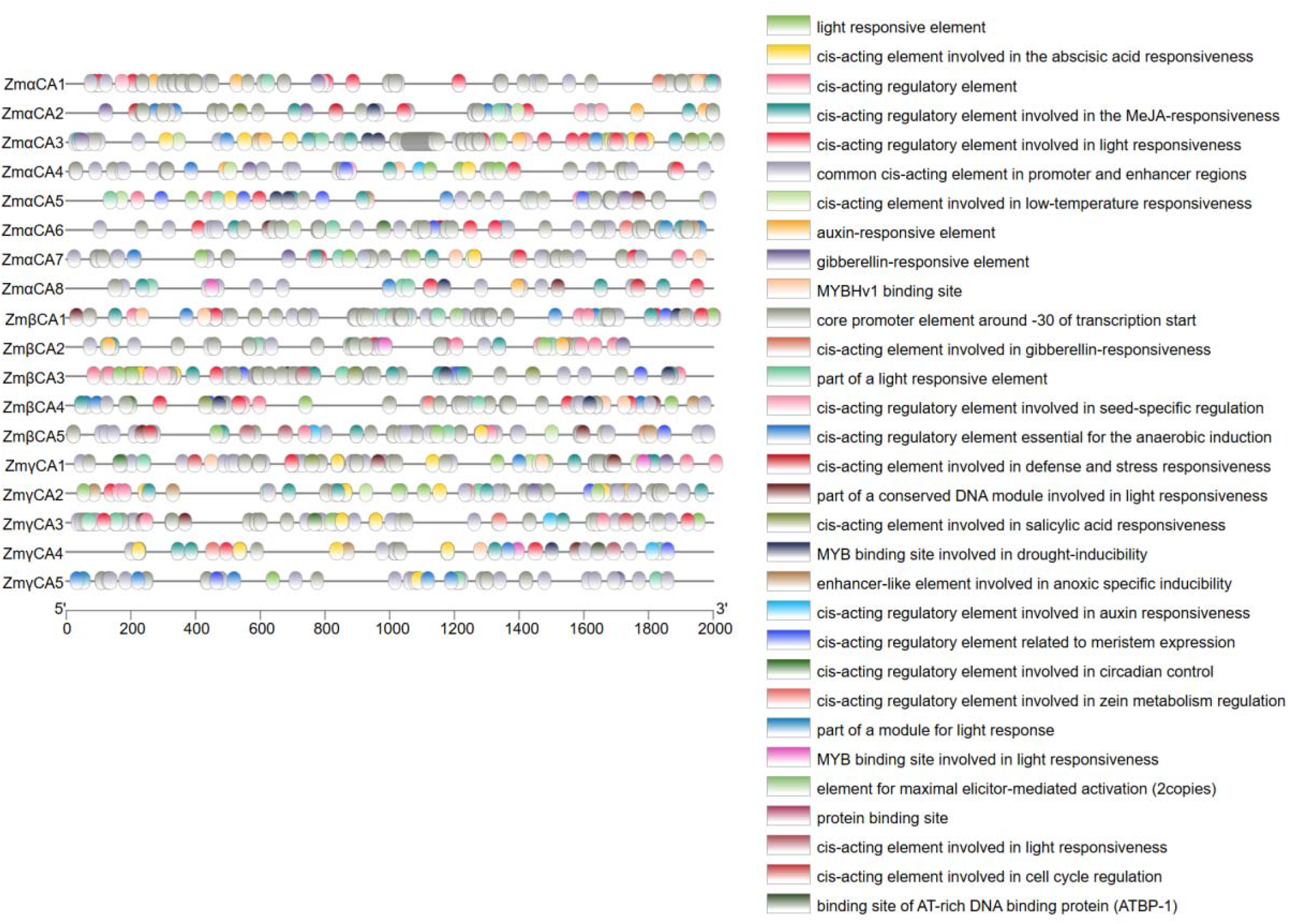
Cis-element analysis of promoters of maize CA family members.

### 2.4 Cis-acting element analysis

ZmCA promoters harbored elements tied to hormones, stress, and development (Raudvere et al., 2019). Using a **hypergeometric test** with **BH FDR** correction (α = 0.05) and a **background of 10**,**000 length-matched random maize promoters** (one per gene), we observe significant enrichment for ABRE (fold=2.1, FDR=0.002), TCA-element (fold=1.8, FDR=0.01), ARE (fold=2.3, FDR=0.001). Subfamily-specific: α-CAs GT1-motif/G-box (light, fold=1.9, FDR=0.005); β-CAs ABRE/ARE (abiotic stress); γ-CAs TCA/LTR (biotic/cold) (Castro-Mondragon et al., 2022).

### 2.5 Synteny analysis of CA genes

Chromosomal segmental duplication, tandem duplication, and WGD are key drivers of gene family expansion in plants, enabling copy number increases, functional diversification, and adaptive innovation (e.g., improved photosynthesis and stress tolerance). Using One-Step MCScanX-superfast in TBtools II, we analyzed within-species synteny of ZmCAs and extended comparisons to rice and Arabidopsis.

Within maize (Fig . 4), we detected a tandem duplication involving *ZmαCA1* and *ZmαCA2* on chromosome 1. This α-specific tandem event likely emerged from a recent local duplication, forming a gene cluster that enhances redundancy and regulatory flexibility (e.g., alternative splicing variants adapted to root-cap pH regulation). Six segmental duplication events involving 10 ZmCAs: four within α (*ZmαCA1*/*ZmαCA5, ZmαCA1*/*ZmαCA6, ZmαCA3*/*ZmαCA7, ZmαCA4*/*ZmαCA7*), one within β (*ZmβCA1*/*ZmβCA4*), and one within γ (*ZmγCA2*/*ZmγCA3*). These span multiple chromosomes (e.g., Chr1 with Chr2, Chr8), suggesting that, following an allotetraploid history and subsequent diploidization, maize retained a subset of duplicates to maintain dosage balance while enabling CA family expansion.

**Figure 4.**
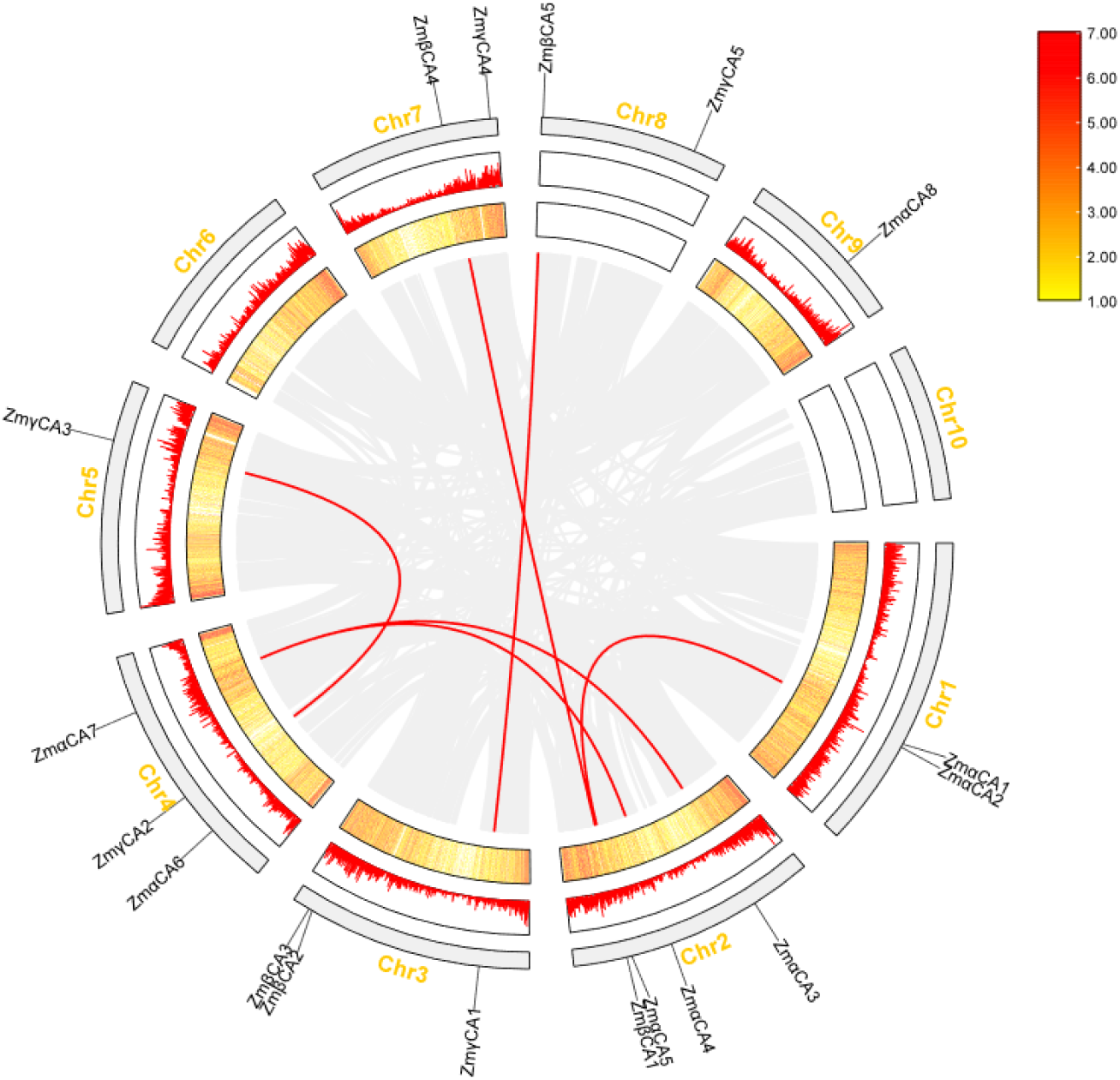
Synteny analysis of the maize CA gene family.

Cross-species collinearity emphasizes conservation: between maize and rice (Fig . 5), 15 orthologous pairs were identified involving nine ZmCAs, predominantly among monocot β and γ orthologs (e.g., *ZmβCA3*–OsβCA), reflecting ancestral presence before the monocot–dicot split and subsequent WGD that reinforced C_4_/C_3_ divergence. In contrast, maize and Arabidopsis share only one direct orthologous pair ((Fig . 5), e.g., *ZmαCA2*–AtαCA, suggesting an ancient origin for α-CAs in land plants and fewer orthologs due to distant relatedness and lineage-specific WGDs. In total, nine ZmCAs have orthologs in both Rice and Arabidopsis, implying pre-divergence conservation with post-divergence positive selection under environmental pressures such as low CO□ or drought. Coupled with promoter elements (e.g., ABRE) and expression patterns (e.g., high *ZmβCA3* in mesophyll), these orthologies point to stress-responsive regulatory networks and potential subfunctionalization contributing to salt tolerance.

**Figure 5.**
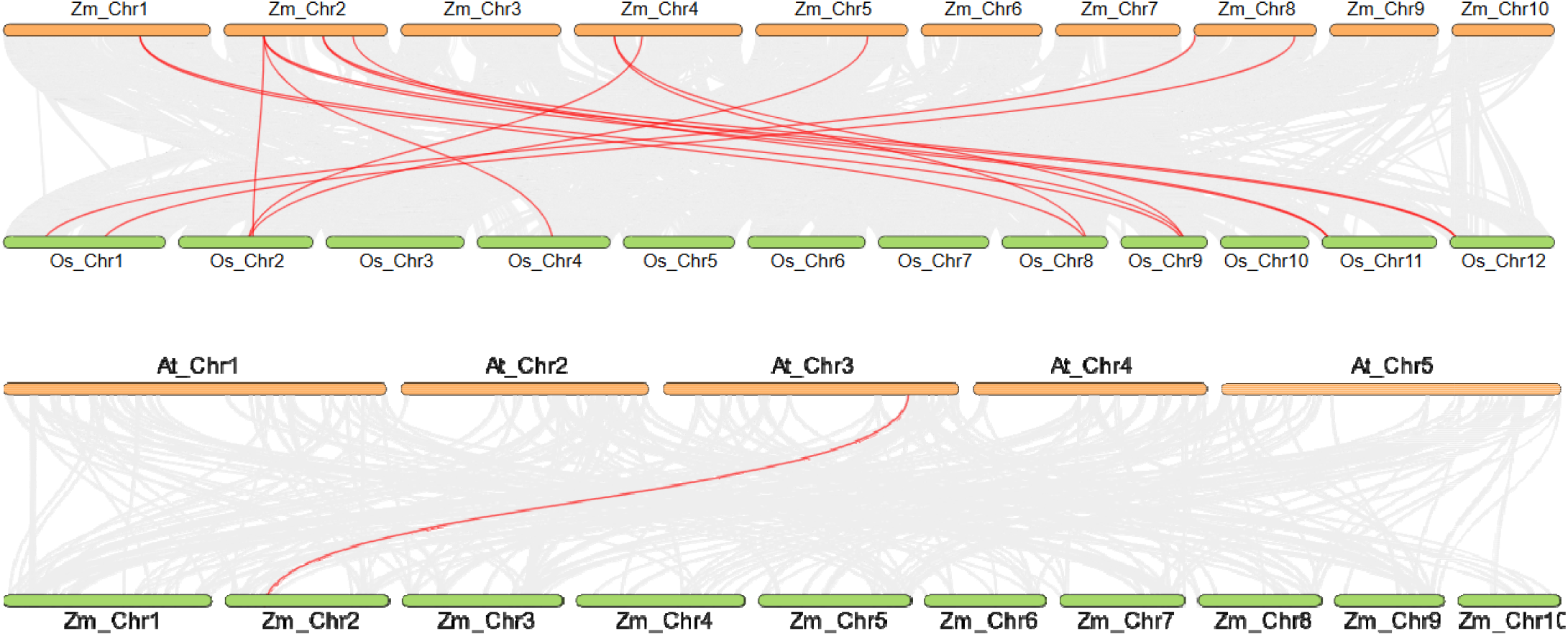
Syntenic relationships of CA genes between Maize, Rice, Arabidopsis. **Note**: Gray lines indicate genome-wide syntenic blocks between species.

These findings deepen our understanding of CA evolutionary dynamics and inform molecular breeding; for example, editing β-CA duplicate pairs to enhance photosynthesis and stress tolerance in maize. (Associated analysis files and results are in: within-species synteny analysis.

### 2.6 Expression analysis of CA family members

RNA-seq across 21 tissues evinced specificity (Figure 6; τ>0.7 for 12 genes) . α-CAs root-predominant (*ZmαCA2* τ=0.82); β-CAs leaf-elevated (*ZmβCA3* log2FC=3.2 vs. root, FDR<1e-5); γ-CAs reproductive. Clustering: ward.D2 on Z-score TPM (Robinson et al., 2010; Murtagh and Legendre, 2014). The analysis reveals tissue specificity and developmental stage dependence, highlighting CA roles in CO□ transport, photosynthesis, pH regulation, and stress responses. In C_4_ maize, high β-CA expression typically correlates with mesophyll-driven carbon concentration, supporting PEPC activity and photosynthetic efficiency under stress (Fig . 6).

**Figure 6.**
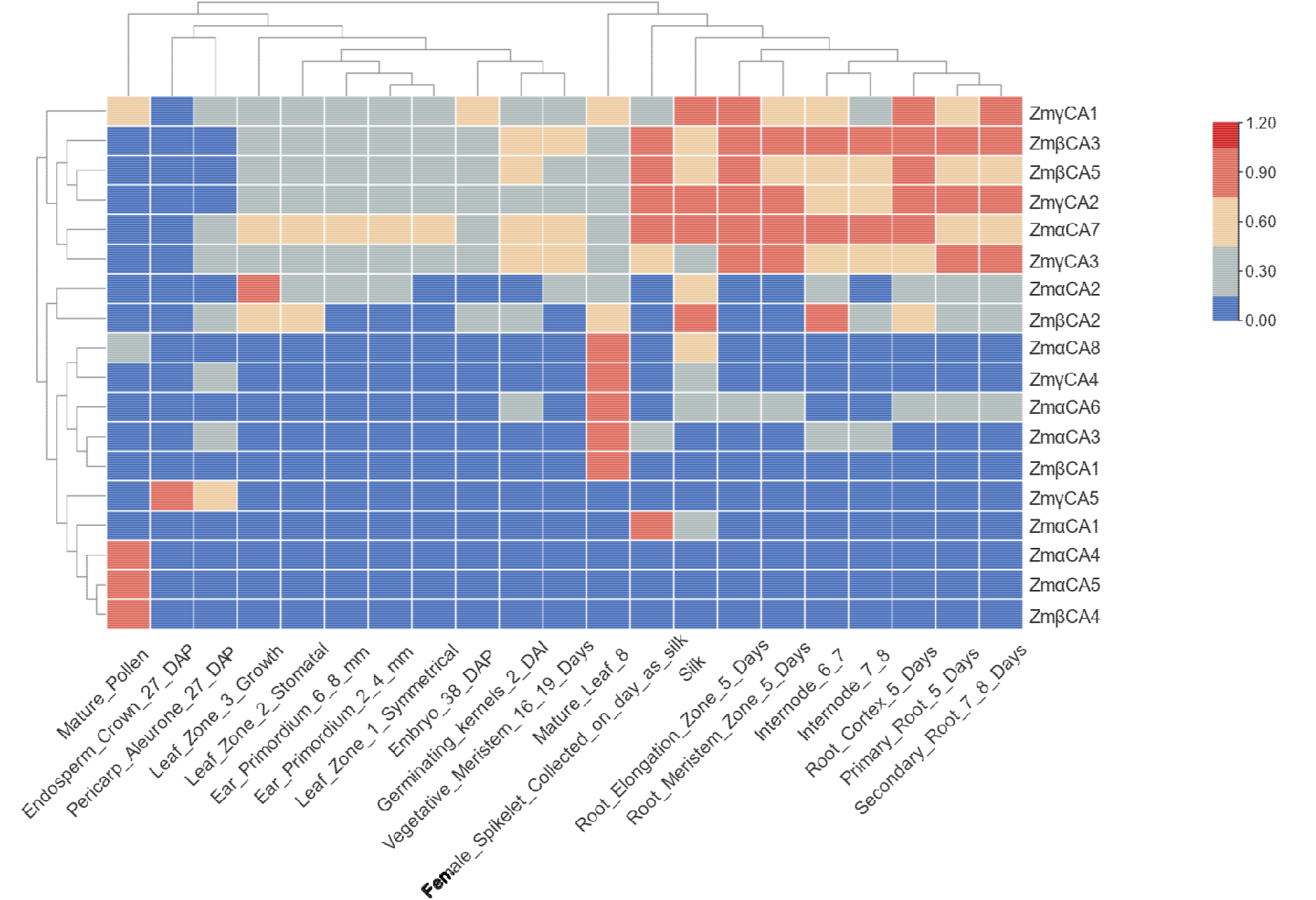
Expression patterns of CA family members across tissues heatmap. **Note**: Z-score TPM (blue low, red high); rows genes, columns tissues; dendrogram ward.D2.

## 3. Single-Cell Transcriptomic Analysis of CA Expression Across Maize Organs

### 3.1 Methods and overall expression overview

Heatmaps across cell types (Fig . 7) unveiled CA subfamily patterns: γ-CAs broadly constitutive, aligning with mitochondrial metabolic roles (DiMario et al., 2017); β- and α-CAs tissue-biased, with β-CAs in photosynthetic cells and α-CAs in supportive niches, evincing partitioning for C_4_ optimization and resilience (Fabre et al., 2024; Engineer et al., 2021). Mesophyll cluster variance (CV >0.5 for β-CAs) suggests stochastic or microenvironmental modulation (Ludwig, 2016).

**Figure 7.**
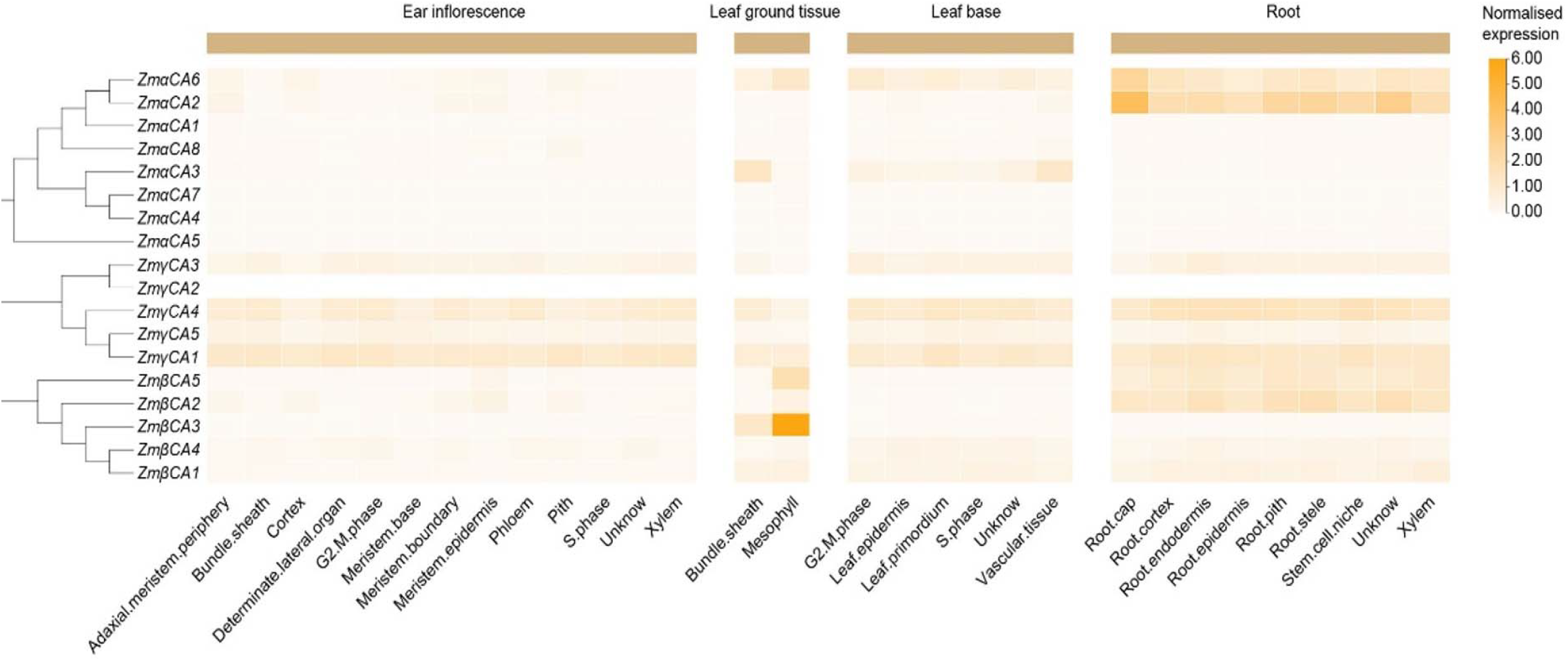
CA family expression across cell types in three organs. **Note**: Heatmap rows are cell types; columns are tissues (ear inflorescence, leaf ground tissue, leaf base, root). Color gradient indicates mean expression (orange high, white low). Left dendrogram shows subfamily clustering.

### 3.2 Expression in ear inflorescences

Ear inflorescences, dynamic hubs for reproductive fate, may leverage CA pH gradients for primordium patterning and signaling (Sun et al., 2024a). γ-CAs predominated, with *ZmγCA1* upregulated (log2FC=1.5, FDR<0.01, pct.1=0.78) versus α/β, implying metabolic stabilization during maturation (Fabre et al., 2024). UMAPs (Fig . 8) depicted diffuse γ-CA distribution across protoderm, meristem, and procambia, lacking subtype enrichment (FDR>0.05) (Sun et al., 2024b). This uniformity contrasts vegetative patterns, echoing rice anthesis CO□ buffering (Hibberd et al., 2019). Subtle *ZmγCA3*/*ZmγCA4* elevations in floral precursors (log2FC=1.2, FDR=0.02) suggest sex determination involvement, potentially via ABA-responsive promoters. This extends maize tassel spatial transcriptomics, positioning γ-CAs as redox/ion balancers amid proliferation.

**Figure 8.**
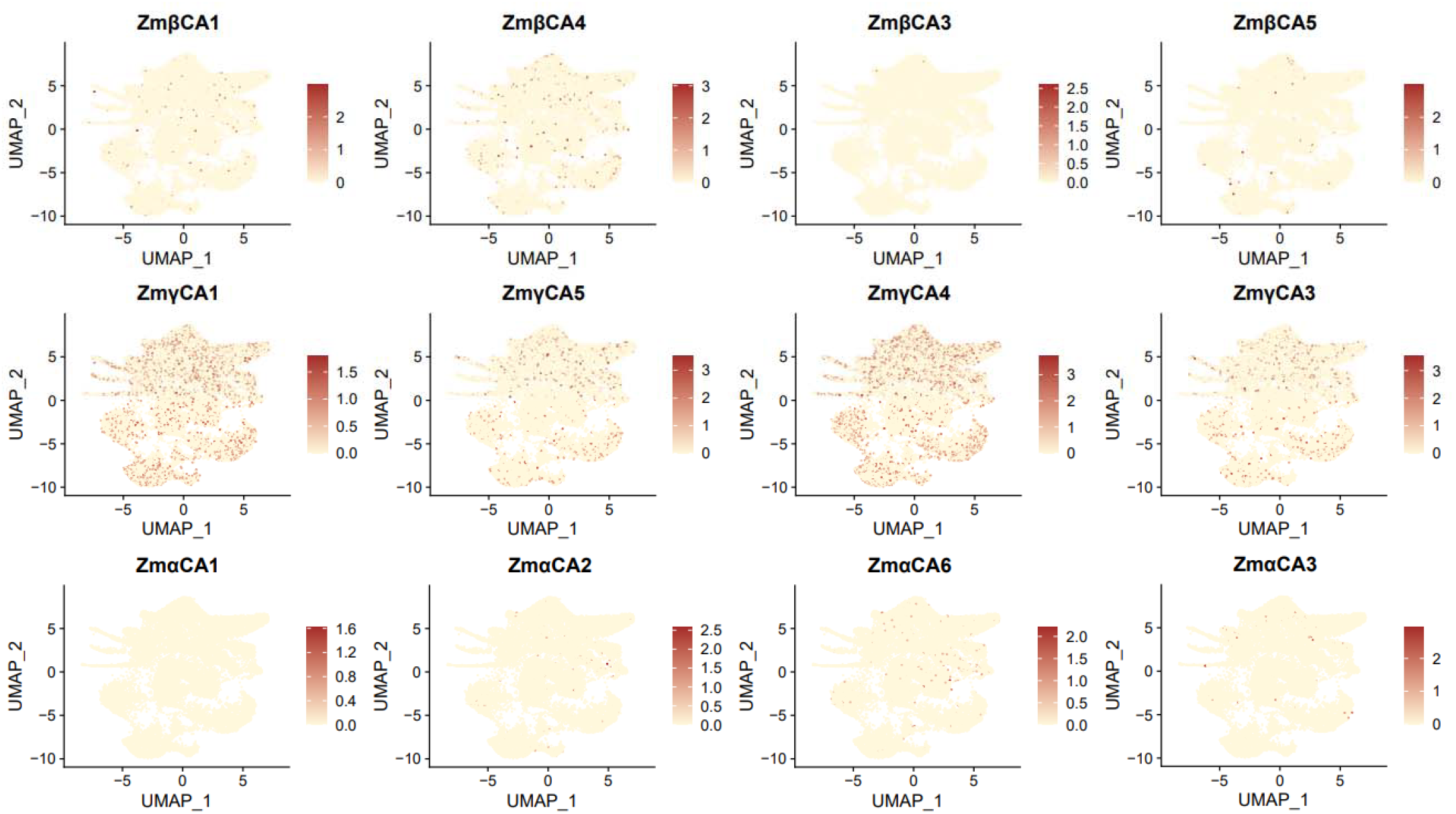
UMAPs of CA family expression in ear inflorescences. **Note**: Each subpanel corresponds to one ZmCA gene; points are single cells; the color gradient (red high, yellow low) denotes expression; UMAP axes indicate cluster structure.

### 3.3 Cell-type-specific expression in leaves

Maize leaves, C_4_ photosynthesis epicenters, compartmentalize architectures where CAs facilitate CO□ mechanisms, mitigating photorespiration under stress. β-CAs enriched in ground tissue, with *ZmβCA3* upregulated in mesophyll (log2FC=2.8, FDR<1e-6, pct.1=0.85). UMAPs (Fig . 9) highlighted *ZmβCA3* foci in mesophyll, correlating with C_4_ markers *ZmPEPC* (r=0.72, p<1e-10), implying light-responsive regulation.

**Figure 9.**
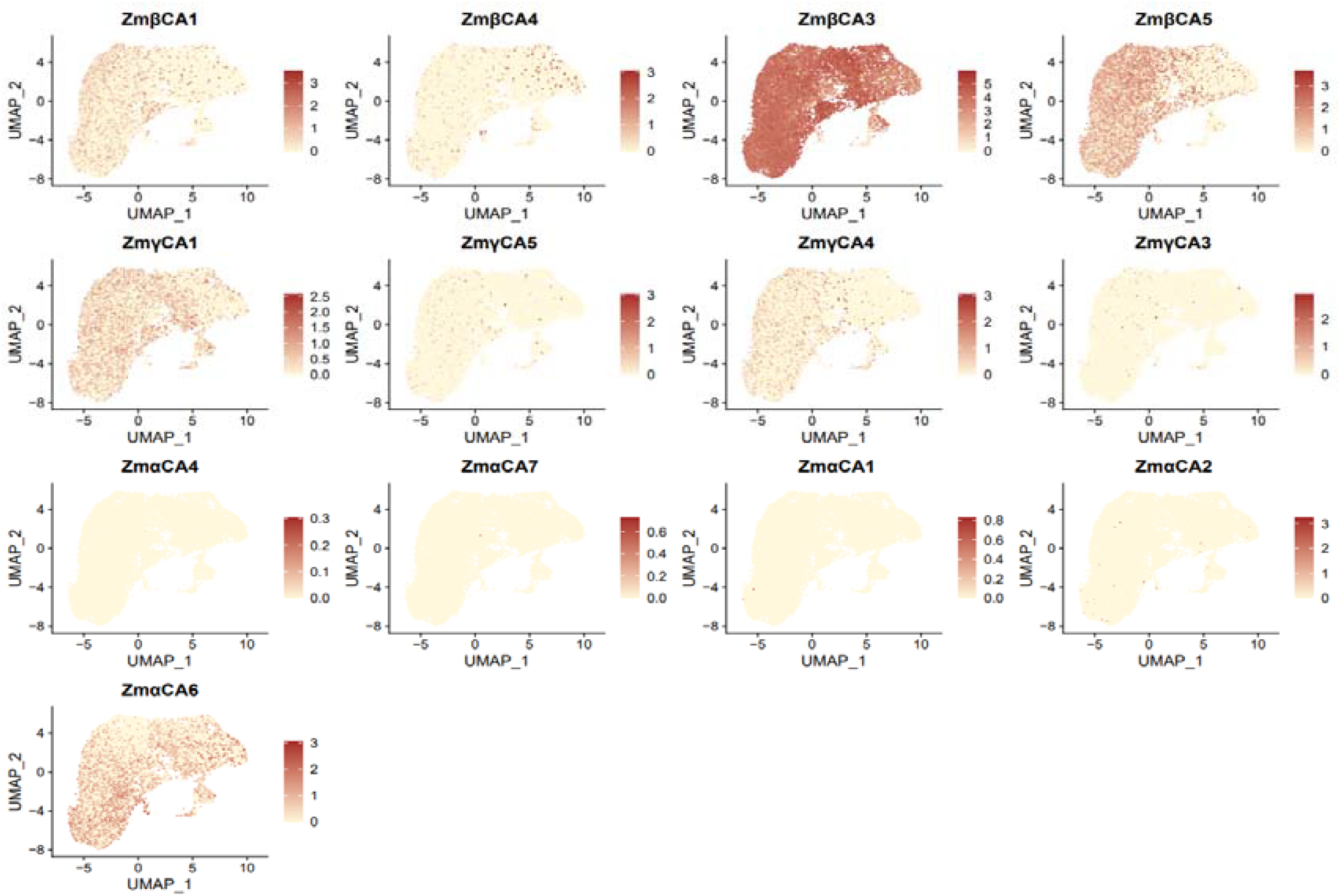
UMAPs of CA family expression in leaf ground tissue.

Conversely, α-CAs prevailed in non-ground tissues, *ZmαCA2* in vascular (log2FC=1.9, FDR<1e-4, pct.1=0.72), aiding pH/ion homeostasis for metabolite flux (Fig . 10). This mirrors sorghum vascular mitigation of hydraulic stress (Engineer et al., 2021). Heterogeneity underscores C_4_ adaptations: β-CAs mesophyll specialization, α-CAs vascular support, priming drought-resilient engineering.

**Figure 10.**
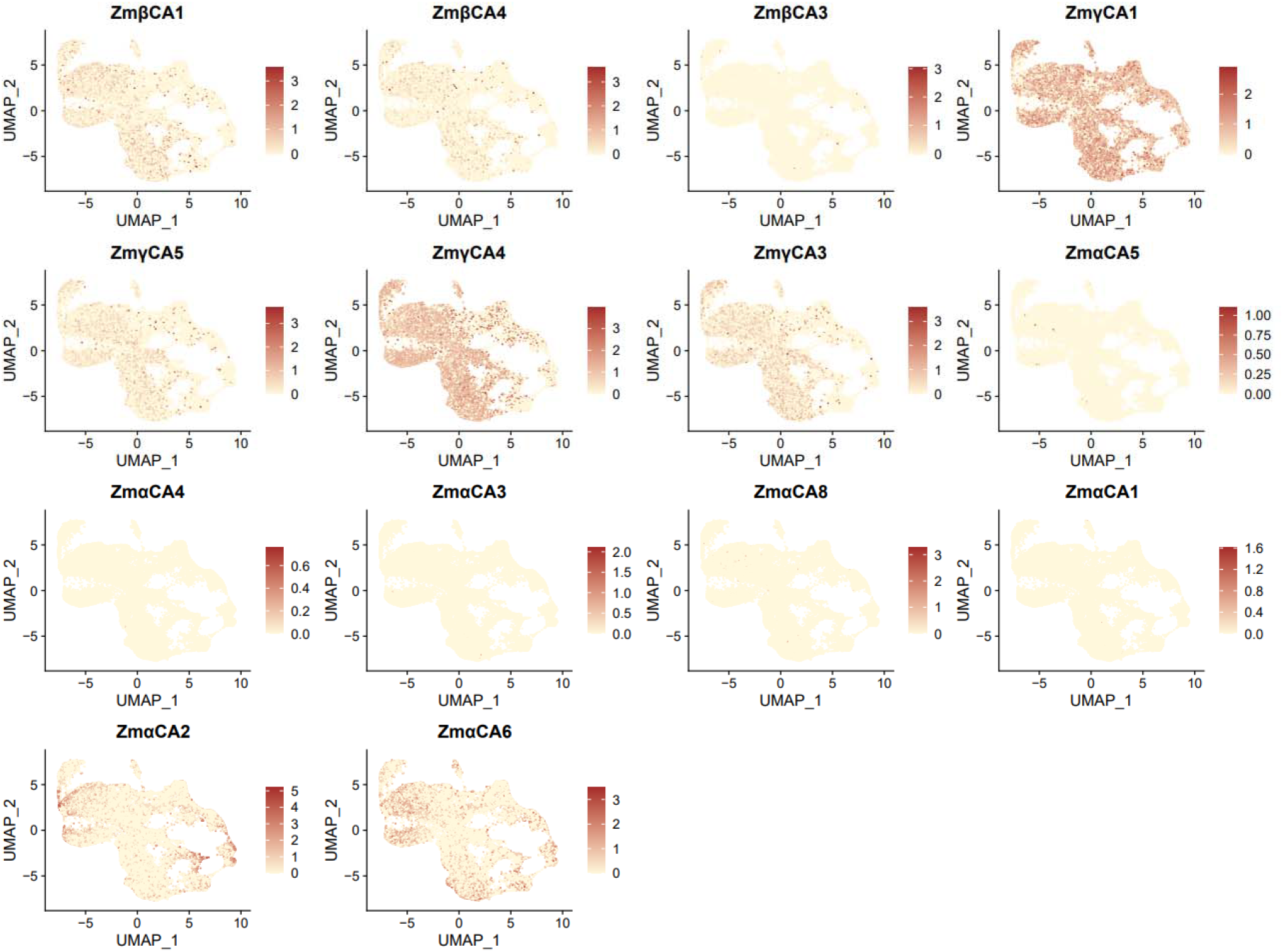
UMAPs of CA family expression in the root apex. **Note**: Each subpanel corresponds to one ZmCA gene; points are single cells; the color gradient (red high, yellow low) denotes expression; UMAP axes indicate cluster structure. Focus on root apex; highlight γ-CA expression across clusters and α-CA expression in the root cap.

### 3.4 Expression patterns in the root apex and functional implications

Root apices sense environmental cues, where CAs may modulate pH/ion fluxes for elongation and stress (Zhang et al., 2025b). γ-CAs dominated, *ZmγCA1* robust across zones (log2FC=1.7, FDR<0.001, pct.1=0.81), likely sustaining respiration amid nutrient hypoxia. UMAPs (Fig . 10) showed uniform γ-CA in meristem / elongation (CV<0.3), indicating housekeeping.

α-CAs like *ZmαCA2* enriched root cap (log2FC=2.1, FDR<1e-5, pct.1=0.76), facilitating gravity/mucilage pH for soil navigation/defense, this parallels Arabidopsis aluminum tolerance. Functionally, γ-CAs buffer hypoxia, α-CAs frontier abiotic responses. Integrating bulk data illuminates networks, e.g., ABA via ABRE, informing root-targeted breeding for water efficiency (Rudenko et al., 2003; Chen et al., 2019).

## 4. Discussion and Conclusions

Our comprehensive delineation of the carbonic anhydrase (CA) gene family in maize clarifies how duplication history, subfamily diversification, subcellular targeting, and regulatory wiring collectively sustain C_4_ performance and stress acclimation. By integrating comparative genomics, synteny, and multi-resolution transcriptomics, we resolve subfamily-specific trajectories and nominate tractable entry points for climate-ready improvement (Fig . 11).

**Figure 11.**
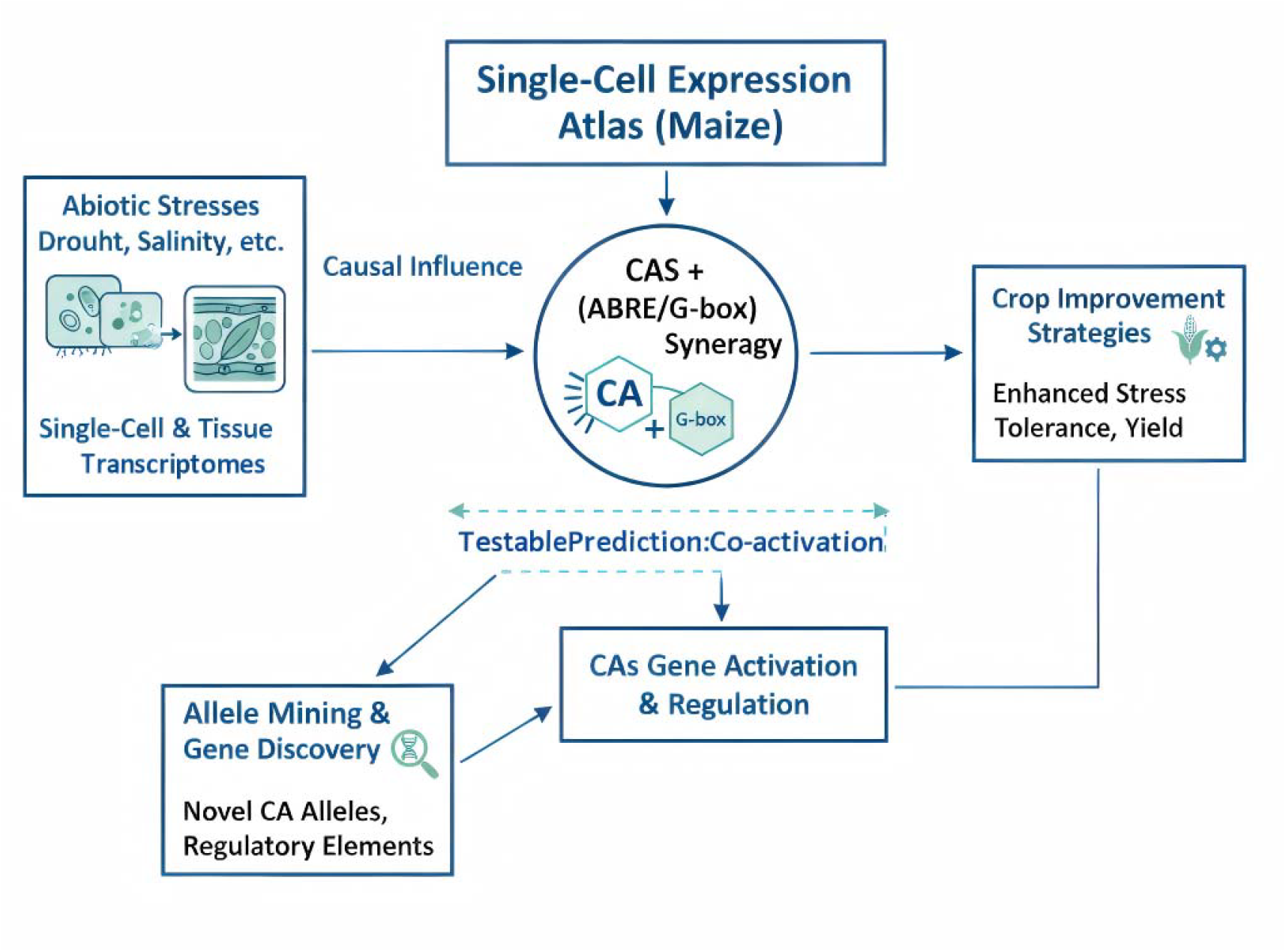
Maize CarbonicAnchyrnaseRegulatory Model: Single-CellAtlasHypthesis. **Note:** This diagram illustrates a workflow where single-cell transcriptomic data from maize under abiotic stress conditions is analyzed to uncover causal influences. The synergy of CAS and ABRE-G-box mechanisms is leveraged for gene activation and regulation, supported by allele mining and gene discovery. These efforts culminate in testable predictions and co-activation strategies, driving the development of improved crop varieties with enhanced stress tolerance and yield.

### 4.1 Identification, Classification, and Evolutionary Conservation

The 18 CA genes we identify (8 α, 5 β, 5 γ) fit the multi-isoform paradigm across angiosperms and mirror prokaryotic origins, yet the maize portfolio is tuned to C4 requirements. The β-CA complement—together with chloroplast targeting and repetitive domains—supports robust bicarbonate provision and proton buffering in chloroplasts under fluctuating CO_2_. Physicochemical breadth (GRAVY −0.49 to 0.142) and thermostability suggest catalytic versatility across cellular niches.

Phylogeny and synteny indicate deep conservation with monocot-specific elaboration: maize shares substantially more orthologs with Rice (15) than Arabidopsis (3), consistent with a 50 Mya monocot–dicot split. Ks peaks and segmental collinearity implicate ancient WGD and segmental duplication in family expansion, while Ka/Ks < 1 indicates pervasive purifying selection preserving core function despite dosage shifts. Importantly, recent discovery of a large deletion in maize β-CA (*cah1*) yielding heightened activity and an environment-specific decoupling of δ13C from water-use efficiency (WUE) and photosynthesis (Twohey et al., 2025) illustrates how adaptive neofunctionalization can arise within an overall purifying backdrop. Taken together, these patterns support a model in which WGD-driven retention and dosage balance maintain enzymatic capacity, while selected β-CA variants tailor CO_2_-concentrating mechanisms (CCMs) to modern stress regimes—a plausible contributor to C_4_ superiority over C_3_ lineages.

### 4.2 Diversity in Gene Structures and Regulatory Elements

Subfamily architectures align with regulatory demands: compact β-CAs favor rapid transcriptional responses to photosynthetic cues; intron-rich α/γ-CAs expand isoform diversity through alternative splicing under stress (DiMario et al., 2023). Motif divergence is matched by organelle targeting—β-CAs harbor chloroplast transit peptides suited for C_4_ compartments, whereas γ-CAs carry mitochondrial import signals, consistent with roles in energy metabolism and ion/pH homeostasis.

Promoter analyses uncover enriched ABREs (fold 2.1, FDR 0.002), salicylic acid–responsive TCA elements (fold 1.8, FDR 0.01), and antioxidant-responsive elements (AREs; fold 2.3, FDR 0.001), embedding CAs within ABA, defense, and redox networks. Co-occurring ABRE–G-box combinations imply synergistic drought–light regulation by *bZIP/ABI3*-like factors. Local tandem clusters (e.g., on chromosome 1) flag co-evolutionary hotspots that could enable coordinated tuning, contrasting with more dispersed arrangements in polyploid wheat (Chen et al., 2023). Convergence with Arabidopsis, where β-CAs and bicarbonate induce *ERF/AP2* modules under hypoxia, drought, and high light (Białas et al., 2024), further suggests a conserved regulatory logic likely co-opted in maize’s ABRE/TCA-rich promoters. These features position CA promoters as tunable modules for stacking multi-stress tolerance.

### 4.3 Expression Patterns and Cellular Heterogeneity

Bulk transcriptomes reveal tissue-biased deployment: leaf-enriched β-CAs (e.g., *ZmβCA3*, log2FC ∼3 vs. root, FDR < 1e−5) align with bicarbonate supply to PEPC; α-CAs predominate in roots, consistent with pH and ion buffering; γ-CAs are elevated in reproductive tissues, matching mitochondrial energy demands. High tissue specificity (τ > 0.7 for most loci) underscores functional partitioning.

Single-cell RNA-seq resolves cell-type logic: *ZmβCA3* is mesophyll-enriched (log2FC ∼2.8, FDR < 1e−6, pct.1 ∼0.85), complementing PEPC and strengthening the CCM; *ZmαCA2* localizes to vascular and cap cells (log2FC ∼1.9, FDR < 1e−4, pct.1 ∼0.72), suggesting a role in ion flux buffering. γ-CAs exhibit more uniform expression across ear cell types (Sun et al., 2024a), consistent with mitochondrial housekeeping during development. Relative to C3 models, maize CAs skew toward photosynthetic and stress-responsive modules, with scRNA-seq hinting at epigenetic and miRNA regulation masked in bulk profiles (Sun et al., 2024b; Tang et al., 2021). Temporal oscillations consistent with diurnal stomatal control and acclimation (Ludwig, 2016) provide an additional layer of coordination.

Stress-resolved single-cell maps of maize roots under heat implicate cortex cells as primary responders, and while CAs were not profiled directly, conserved response modules support analogous roles for root α-CAs in ion and pH homeostasis under thermal perturbation (Wang et al., 2025). Comparative insights from single-cell C_4_ models (e.g., Bienertia) pointing to plasma-membrane β-CAs that enhance CO_2_ capture motivate spatial transcriptomics and proteomics in maize to pinpoint subcellular localization refinements (Nguyen et al., 2025).

### 4.4 Functional Implications for Stress Responses and Molecular Breeding

Collectively, the CA family forms a distributed hub for stress mitigation across compartments. β-CAs stabilize CCMs under drought and heat by sustaining bicarbonate supply and proton buffering; α- and γ-CAs modulate pH and bicarbonate-linked signaling to maintain cellular homeostasis under salinity and hypoxia (DiMario et al., 2017; Fabre et al., 2024). Promoter ABRE/TCA enrichment mechanistically links CAs to ABA-driven stomatal control, osmotic adjustment, and SA-mediated defense, with AREs integrating redox crosstalk.

Duplication under purifying selection has fostered sub- and neofunctionalization, enabling context-specific deployment that likely contributes to maize’s edge over C_3_ crops. These insights translate into concrete strategies: CRISPR/Cas-based activation or promoter engineering of mesophyll-specific β-CAs (e.g., *ZmβCA3*) to fortify CCMs; marker-assisted selection focused on tandem clusters on chromosome 1; and allele mining for variants predicted to enhance thermostability or chloroplast targeting. Cross-species validation is encouraging—overexpressing C_4_-derived β-CA in Arabidopsis improves photosynthesis, amino acid synthesis, biomass, and WUE (Kandoi et al., 2022)—underscoring the portability of CA-based gains.

Key gaps and tractable next steps remain. Post-transcriptional layers (protein abundance, localization, pH microenvironments) require quantitative proteomics, metabolomics, and in vivo reporters. Isoform-resolved perturbations (CRISPR editing) combined with fluxomics (^13^CO_2_ tracing) can test causality between β-CA dosage and PEPC carboxylation under high VPD and heat. Spatial transcriptomics and proximity labeling could resolve compartment-specific interactomes, while multiplexed promoter swaps or ABRE–G-box edits would probe how motif architecture governs drought–light synergy.

## Conclusions

Maize CAs calibrate inorganic carbon supply, proton flux, and hormone/redox signaling across organelles and cell types to stabilize C_4_ physiology under environmental variability. Evolutionary retention under purifying selection has preserved core catalysis, while selective innovation—especially within β-CAs—has fine-tuned CCM robustness. Exploiting this architecture through precise regulatory and subcellular engineering offers a credible path to climate resilience and sustainable yield gains.

This study elucidates the structural, evolutionary, and expressional facets of the maize CA gene family, revealing their pivotal roles in C_4_ photosynthesis and multifaceted stress responses. By integrating genome-wide identification, phylogenetic conservation, regulatory diversity, and single-cell resolution expression, we establish a foundational framework for understanding CA contributions to maize adaptation. These insights not only deepen our appreciation of C_4_ evolutionary innovations but also pave the way for targeted molecular breeding strategies, such as CRISPR-enhanced resilience and MAS for yield optimization. Future investigations incorporating proteomics, metabolomics, and transgenic approaches will further unravel dynamic interactions, ultimately supporting the development of climate-smart maize varieties.

## Materials and Methods

### Identification of CA family members in maize and physicochemical property analysis

Genomic resources for Zea mays (B73 RefGen_v5), Oryza sativa (MSU7), and Arabidopsis thaliana (TAIR10) were sourced from respective repositories (Jiao et al., 2017). Arabidopsis CA sequences informed BLASTP queries (e-value ≤ 1e-5, coverage ≥ 50%, identity ≥ 30%) for maize homologs (Altschul et al., 1990). Concurrently, Pfam v36 profiles (α-CA: PF00194; β-CA: PF00484; γ-CA: PF00132) guided HMMER v3.3.2 searches (--cut_ga) (Finn et al., 2016; Eddy, 2011). Intersectong candidates underwent redundancy removal via CD-HIT (90% identity) (Li and Godzik, 2006). Domains were corroborated with Pfam v36, NCBI Batch CD-Search, and SMART v9 (Marchler-Bauer et al., 2015; Letunic and Bork, 2018). Manual curation ensured CA domain integrity. ProtParam derived physicochemical traits (Gasteiger et al., 2003), and subcellular predictions stemmed from TargetP 2.0, ChloroP, and MitoFates consensus (confidence ≥0.5) (Emanuelsson et al., 2000; Small et al., 2004).

### Gene structure and conserved motif analysis of maize CA genes

Exon-intron configurations were extracted from maize GFF and genome files (Pertea et al., 2015). MEME v5.5.5 discerned conserved motifs (-nmotifs 10, -minw 6, -maxw 50, ZOOPS, E-value <1e-5) (Bailey et al., 2015). Domains were scrutinized via NCBI Batch CD-Search (Marchler-Bauer et al., 2017). Visualizations employed TBtools II v2.0 (Chen et al., 2023). Motifs aligned to catalytic residues (e.g., Zn-binding His/Cys/Asp) per PROSITE and literature (Sigrist et al., 2013; Rowlett, 2010).

### Phylogenetic analysis of CA proteins

CA sequences from maize, Arabidopsis, and rice underwent MAFFT v7 (L-INS-i) alignment (Katoh and Standley, 2013) and trimAl (-automated1) refinement (Capella-Gutiérrez et al., 2009). IQ-TREE v2 constructed ML phylogeny with ModelFinder-selected LG+F+I+G4, 1000 UFboot, and SH-aLRT supports (Nguyen et al., 2015; Hoang et al., 2018). Rooting at γ-clade reflected prokaryotic origins (Rowlett, 2010).

### Chromosomal mapping, gene duplication, and synteny analysis

ZmCA loci were mapped from GFF using TBtools II v2.0 (Chen et al., 2020). Synteny across maize, Arabidopsis, and rice utilized One-Step MCScanX-superfast (match_size=5, gap_penalty=3, e-value=1e-5) (Wang et al., 2012). Tandems defined as <200 kb with ≤1 intervening gene (Cannon et al., 2010). Orthologs filtered by bidirectional best-hit + synteny (Ostlund et al., 2010). Ka/Ks computed via KaKs_Calculator v2.0 (yn00) (Wang et al., 2010); Ks peaks timed WGD (clock 6.5e-9) (Blanc and Wolfe, 2004).

### Cis-acting element analysis of maize CA genes

Promoter sequences (2,000 bp upstream ATG) were extracted via TBtools II v2.0 and analyzed with PlantCARE and JASPAR Plants (Lescot et al., 2002; Castro-Mondragon et al., 2022). Enrichment employed hypergeometric test (FDR<0.05, BH) (Raudvere et al., 2019). Position and co-occurrence (e.g., ABRE+G-box) were examined in R v4.4 (R Core Team, 2024). Visualizations via TBtools II v2.0 (Chen et al., 2023).

### Gene expression analysis

RNA-seq across 21 tissues/stages sourced from qTeller (Woodhouse et al., 2021; TPM, merged isoforms, ComBat-corrected). Specificity via τ index (>0.7 specific) (Yanai et al., 2005). Clustering: ward.D2 on Euclidean (Murtagh and Legendre, 2014). Differential expression for select genes via edgeR (FDR<0.05) (Robinson et al., 2010).

### Single-cell analysis

scRNA-seq from scPlantDB: ear inflorescences (SRP272727 n=4567; SRP272723 n=3789; SRP272726 n=4123), leaves (SRP281914 n=5678), root apex (SRP145013 n=4892) (Xu et al., 2021; Ortiz-Ramírez et al., 2021). Processed in R v4.4 with Seurat v5.1.0 (Hao et al., 2021): CreateSeuratObject (min.cells=3, min.features=200) (Butler et al., 2018), QC (mt%<20%, rb%<5%), DoubletFinder (pK=0.09) (McGinnis et al., 2019), Harmony integration (dims=1:30) (Korsunsky et al., 2019), SCTransform (vars.to.regress=“percent.mt”) (Hafemeister and Satija, 2019), FindNeighbors (dims=1:30, Elbow >80% variance) (Becht et al., 2019), FindClusters (resolution=0.8, silhouette >0.6) (Traag et al., 2019). Annotations via markers (e.g., mesophyll: ZmPEPC, ZmRBCS; vascular: ZmSWEET13, ZmPPDK; root cap: ZmNAC7) (Nelms and Walbot, 2019; Satterlee et al., 2020) and SingleR against maize atlas (Macosko et al., 2015). Differential expression: Wilcoxon (min.pct=0.1, |log2FC|≥0.25, BH-FDR<0.05) (Wilcoxon, 1945). Heatmaps (Z-score, ward.D2) via ComplexHeatmap v2.20.0 (Gu et al., 2016). UMAPs via ggplot2 v3.5.1 (DimPlot, FeaturePlot) (Wickham, 2016).

### Consent for publication

This article was jointly written by the authors Yonggang Gao, Cheng Zhao,and each author agreed to publish this article. We would like to thank each author for their hard work.

## Additional information

The authors declare that there was no funding for this work.

## Author contributions

Cheng Zhao assumed responsibility for content planning and primary chapter composition within the article. Yonggang Gao was tasked with Figure preparation, experimental design, data analysis, etc., .

